# A toolbox of anti-mouse and rabbit IgG secondary nanobodies

**DOI:** 10.1101/209742

**Authors:** Tino Pleiner, Mark Bates, Dirk Görlich

**Author notes:** Correspondence should be addressed to D.G.; M.B. or T.P.

## Abstract

Polyclonal anti-IgG secondary antibodies are essential tools for many molecular biology techniques and diagnostic tests. Their animal-based production is, however, a major ethical problem. Here, we introduce a sustainable alternative, namely nanobodies against all mouse IgG subclasses and rabbit IgG. They can be produced at large scale in *E. coli* and could thus make secondary antibody-production in animals obsolete. Their recombinant nature allows fusion with affinity tags or reporter enzymes as well as efficient maleimide chemistry for fluorophore-coupling. We demonstrate their superior performance in Western Blotting, both in peroxidase- and fluorophore-linked form. Their site-specific labeling with multiple fluorophores creates bright imaging reagents for confocal and super-resolution microscopy with much smaller label displacement than traditional secondary antibodies. They also enable simpler and faster immunostaining protocols and even allow multi-target localization with primary IgGs from the same species and of the same class.

## Introduction

Mouse and rabbit antibodies are fundamental tools for numerous basic research techniques as well as medical diagnostic assays. The detection or immobilization of these primary antibodies is most often performed indirectly via polyclonal anti-IgG secondary antibodies. Yet, the need for a continuous supply of anti-IgG sera requires keeping, immunizing, bleeding and eventually sacrificing large numbers of goats, sheep, rabbits, or donkeys, which is not only costly but also a major animal welfare and ethical problem (Shen, 2013; Reardon, 2016). Furthermore, every new batch of serum contains another heterogeneous mixture of antibodies, which need to be affinity-purified on IgG columns and then depleted (by pre-adsorption) of nonspecific and crossreacting antibodies. Moreover, the success of this procedure has to be laboriously quality controlled each time. The large size of secondary antibodies (~10-15 nm; 150 kDa) is also a disadvantage, since it limits tissue penetration and introduces a considerable label displacement, reducing the obtainable image resolution by super-resolution fluorescence microscopy methods (Ries et al., 2012; Szymborska et al., 2013; Pleiner et al., 2015). Their non-recombinant nature further precludes genetic engineering i.e. tagging or fusion to reporter enzymes.

Why then, have recombinant anti-IgG detection reagents not yet replaced polyclonal secondary antibodies? The major issue is regarding signal strength. The signal in traditional immunofluorescence, for example, is amplified by: (i) multiple secondary IgG molecules binding to distinct epitopes of a primary antibody; (ii) a large IgG tolerating many labels per molecule; and (iii) by their bivalent binding mode exploiting avidity for high affinity target recognition. In the light of these facts, it appears very challenging to achieve comparable signal levels with a small, monovalent and monoclonal reagent.

Yet, we considered nanobodies, single-domain antibodies derived from camelid heavy-chain antibodies (Hamers-Casterman et al., 1993; Arbabi Ghahroudi et al., 1997; Muyldermans, 2013), as perhaps the best candidates for such reagents. Due to their small size (~3×4 nm; 13 kDa), the possibility of their renewable production as recombinant fusion proteins, as well as favorable biophysical properties, nanobodies attracted considerable attention as powerful tools in cell biology (Helma et al., 2015), structural biology (Desmyter et al., 2015) and as future therapeutic agents (Van Bockstaele et al., 2009; Kijanka et al., 2015). They are particularly useful for super-resolution imaging (Ries et al., 2012; Szymborska et al., 2013; Pleiner et al., 2015; Göttfert et al., 2017; Traenkle and Rothbauer, 2017). The resolving power of some of the best microscopes reported to date (e.g. ~6 nm by Balzarotti et al., 2017; ~10-20 nm by Huang et al., 2016 or Xu et al., 2012) may be reduced due to the offset between fluorescent label and target introduced by primary and secondary antibodies (20-30 nm). Site-specifically labeled nanobodies represent a promising solution to this problem, since they can place fluorophores closer than 2 nm to their antigen and, despite their small size, even tolerate up to three dyes (Pleiner et al., 2015).

In this study, we describe the generation of a comprehensive toolbox of nanobodies against all mouse IgG subclasses and rabbit IgG. This work required very extensive optimizations of our routine nanobody selection efforts, such as a time-stretched and thus affinity-enhancing immunization scheme, subsequent affinity maturation including off-rate selections, as well as testing and improving ~200 initial candidates. When labeled site-specifically with fluorophores, the resulting nanobodies performed remarkably well in Western Blotting and immunofluorescence. In contrast to polyclonal secondary antibodies, they even allow a single-step multicolor labeling and co-localization. In Stochastic Optical Reconstruction Microscopy (STORM) of microtubules, an anti-mouse kappa light chain nanobody showed greatly reduced fluorophore offset distances, suggesting its use as a superior alternative to traditional anti-mouse secondary antibodies. Moreover, we show that anti-IgG nanobodies can be conjugated to horseradish peroxidase (HRP) or expressed as fusions to ascorbate peroxidase (APEX2) (Lam et al., 2015) and thus used for enhanced chemiluminescence Western blotting or colorimetric ELISAs or immuno-EM detection. These monoclonal recombinant nanobodies are thus perfect substitutes for conventional animal-derived polyclonal secondary antibodies. We envision that they can be engineered to enable a more versatile use of the plethora of existing antibodies and even allow the development of more sophisticated antibody-based diagnostic tests.

## Results

### A comprehensive anti-IgG nanobody toolbox

We immunized two alpacas separately with polyclonal mouse or rabbit IgG and used chemically biotinylated mouse monoclonal antibodies (mAbs) of defined subclasses as well as rabbit IgGs for phage display selections of nanobodies from the resulting immune libraries. First results with the initially obtained anti-IgG nanobodies were rather disappointing, i.e. we experienced dim and noisy signals in immunofluorescence as well as in Western blots. We reasoned that an increase in affinity and specificity might yield improved reagents and therefore re-immunized the animals after a one-year pause. For this, we used IgGs pre-bound to multivalent particulate antigens expected to provide strong T-helper cell epitopes. Moreover, we increased the stringency of the subsequent phage display selections by lowering the bait concentration down to the femtomolar range, which should not only select *per se* for sub-nanomolar binders, but also bring displayed nanobodies in direct competition with each other, because the number of bait molecules was up to 1000-fold lower than the number of displaying phages. Finally, we performed *in vitro* affinity maturations by random mutagenesis and further rounds of phage display, this time also combined with off-rate selections. In this way, we obtained a large toolkit of anti-rabbit and anti-mouse IgG nanobodies (Fig. 1 A).

**Figure 1.**
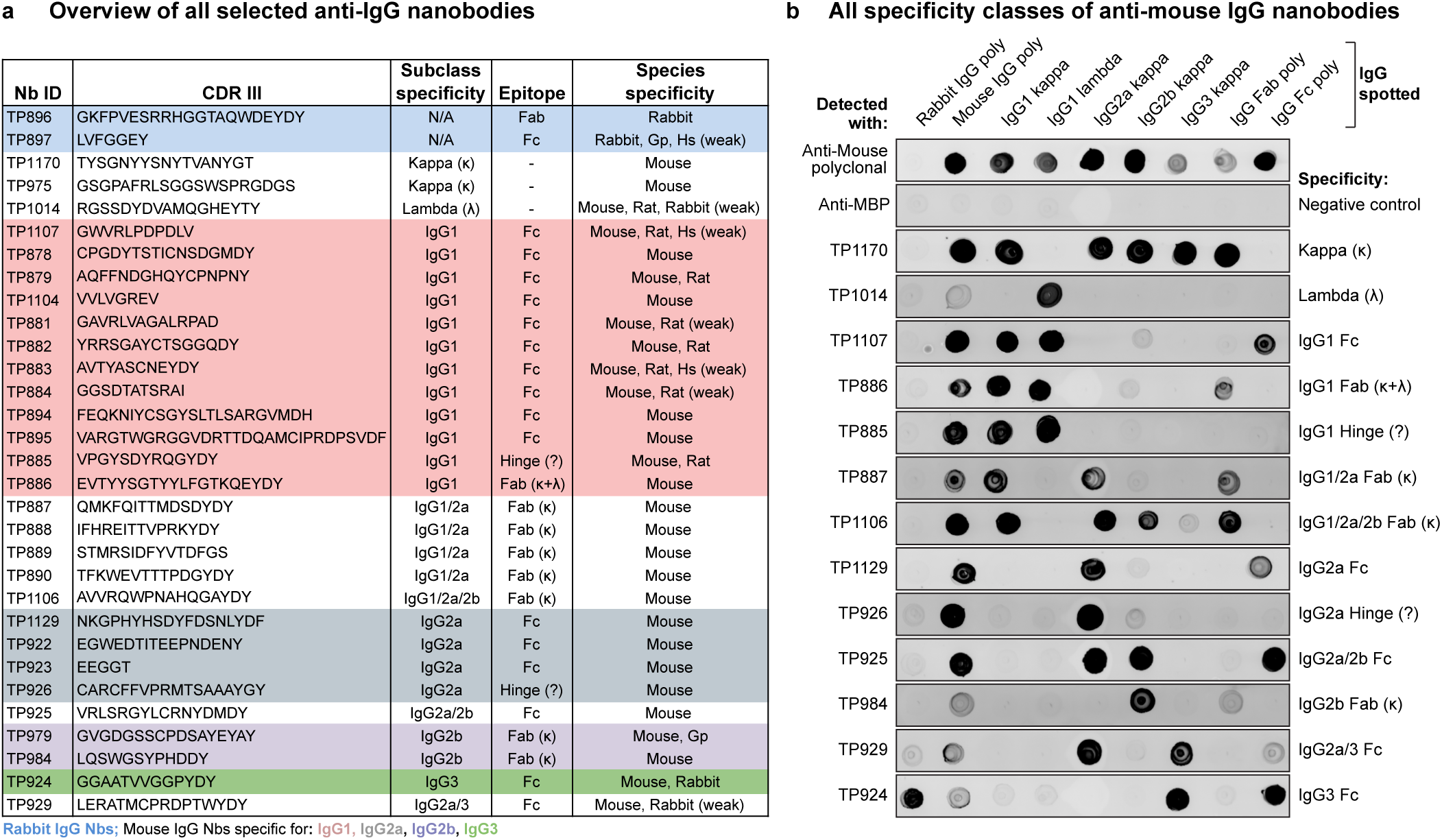
Characterization of the anti-IgG nanobody toolbox. **(A)** Overview of all identified anti-IgG nanobodies. The obtained nanobodies were characterized for IgG subclass specificity, epitope location on Fab or Fc fragment and species crossreactivity (Fig. S1 A). The protein sequences of all anti-IgG nanobodies can be found in Supplementary Table 1. Nb = nanobody; CDR III = Complementarity-determining region III; Gp = Guinea pig; Hs = Human; κ = kappa light chain; λ = lambda light chain; Fab = Fragment antigen-binding, Fc = Fragment crystallizable. **(B)** IgG subclass reactivity profiling of selected anti-mouse IgG nanobodies representing all identified specificity groups. The indicated IgG species were spotted on nitrocellulose strips and the strips blocked with 4 % (w/v) milk in 1x PBS. Then 300 nM of the indicated tagged nanobodies were added in milk. After washing with 1x PBS, bound nanobodies were detected using a fluorescent scanner. Note that the signal strength on poylclonal IgG depends on the relative abundance of the specific subclass (e.g. IgG2b and IgG3 are low-abundant) or light chain (kappa : lambda ratio = 99:1). TP885 and TP926 showed no detectable binding to polyclonal Fab or Fc fragment and might bind to the hinge region. MBP = maltose binding protein; poly = polyclonal.

All nanobodies were extensively characterized for subclass specificity, epitope location on Fab or Fc fragment and crossreactivity to IgGs from other species (Fig. 1 B and Fig. S1 A). Their full protein sequences are listed in Supplementary Table 1, and clones are available on request and will also be distributed through Addgene. Notably, we identified nanobodies against all four mouse IgG subclasses and the sole rabbit IgG subclass. Strikingly, many anti-mouse IgG nanobodies target IgG1, which represents the most abundant subclass of commercially available mouse mAbs (~62-64 %), followed by IgG2a (~22-24 %) and the less frequent IgG2b (~13 %) and IgG3 (~1-2 %). Since the vast majority (~99 %) of mouse mAbs possess a kappa light chain, anti-kappa chain nanobodies promised to be the most broadly useful tools and we therefore actively selected for such binders by swapping the IgG heavy chain subclass during sequential selection rounds. For the identification of binders targeting the rare lambda chain, we had to pre-deplete the nanobody immune library of heavy chain and kappa chain-binders. Some of the identified nanobodies have mixed specificities, e.g. multiple mouse Fab-binders target an interface between kappa light chain and IgG1 or IgG2a heavy chain. Most anti-mouse IgG nanobodies are exclusively mouse-specific, while others additionally crossreact with rat IgG (Fig. S1 A). The anti-rabbit IgG nanobody TP897 also efficiently recognizes guinea pig IgG. All nanobodies were produced by cytoplasmic expression in *E. coli*, mostly with an N-terminal His-NEDD8-tag for purification by Ni(II) chelate affinity capture and proteolytic release (Frey and Görlich, 2014). They were further equipped with ectopic cysteines for subsequent maleimide labeling reactions (Pleiner et al., 2015). Without further optimization, we typically obtained yields of 15 mg per liter of bacterial culture, which already suffices for a million immunofluorescence stains or 200 liters of Western blotting solution (see below).

We first assessed if the anti-IgG nanobodies were specific and could purify their IgG target from its common source. Anti-rabbit IgG nanobodies TP896 and TP897 isolated polyclonal rabbit IgG from crude rabbit serum with high specificity (Fig. S1 B). Likewise, anti-mouse IgG nanobodies TP881 and TP885 could purify an IgG1 mAb from hybridoma cell culture supernatant (Fig. S1 C). Notably, nanobody-bound IgG was released under physiological conditions using SUMOStar protease cleavage (Pleiner et al., 2015). The main virtue of this approach is perhaps not to purify IgGs from sera, but to perform immune-affinity purifications of antigens or antigen complexes that have been pre-bound to the primary antibodies. In contrast to traditional IPs, this approach allows to release the purified complexes under fully native conditions.

### Western blotting with horseradish peroxidase-conjugated anti-IgG nanobodies

We next tested the performance of anti-IgG nanobodies as detection reagents in Western Blotting, which is a major application for secondary antibodies. A popular mode of signal detection in Western Blotting is enhanced chemiluminescence (ECL) in which antibody-horseradish peroxidase (HRP) conjugates are used. HRP is a heme-containing enzyme that catalyzes the oxidation of luminol in the presence of H_2_O_2_ to yield bright chemiluminescence, which is greatly increased by phenol-derived enhancers. We conjugated maleimide-activated HRP to anti-mouse IgG1 Fc nanobody TP1107 via a C-terminal cysteine (Fig. S2 A) and used the resulting conjugate in ECL Western Blotting. The nanobody-HRP conjugate is functional and outperformed a polyclonal secondary antibody-HRP conjugate from a commercial supplier (Fig. 2 A). The anti-rabbit IgG nanobody TP897 could also be linked to HRP and the resulting conjugate was functional and specific.

**Figure 2.**
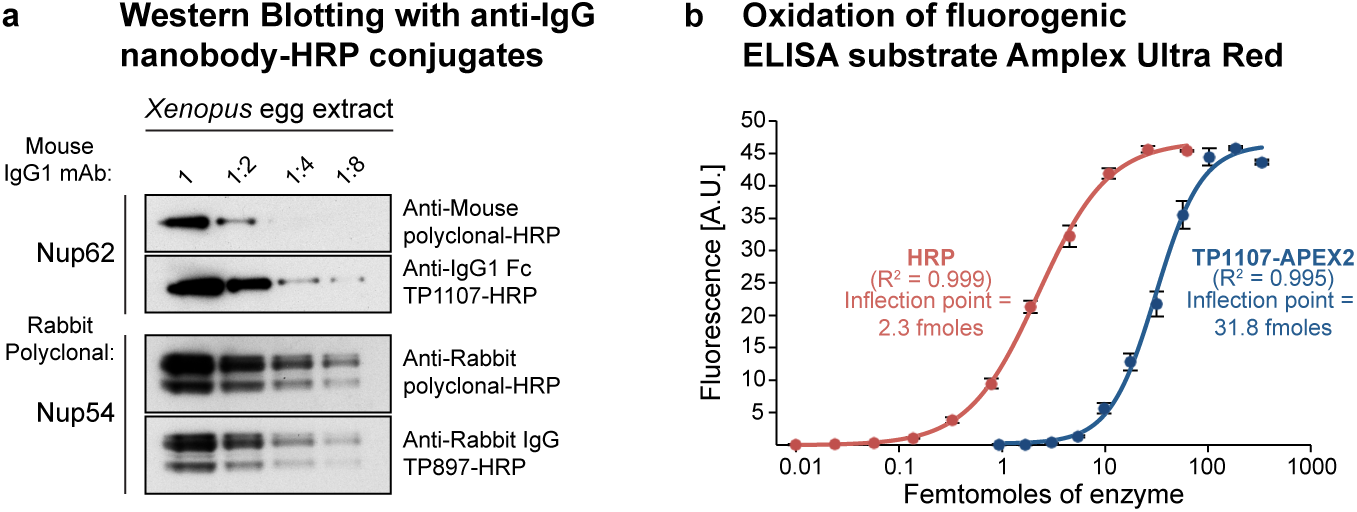
Application of peroxidase-linked anti-IgG nanobodies. **(A)** A twofold dilution series of *Xenopus laevis* egg extract was blotted and probed with anti-Nup62 mouse IgG1 mAb A225. It was then decorated with horseradish peroxidase (HRP)-conjugated goat anti-mouse polyclonal IgG (5 nM) or anti-mouse IgG1 Fc nanobody TP1107 (5 nM) and detected via enhanced chemiluminescence (ECL). Similarly, a rabbit polyclonal antibody targeting Nup54 was decorated with HRP-conjugated goat anti-rabbit polyclonal IgG or anti-rabbit IgG nanobody TP897 (5 nM). **(B)** Oxidation of the fluorogenic ELISA substrate Amplex Ultra Red. A dilution series of pure HRP or recombinant anti-mouse IgG1 Fc nanobody TP1107-Ascorbate peroxidase (APEX2) fusion was incubated with Amplex Ultra Red and H_2_O_2_. Oxidation leads to formation of the fluorescent compound resorufin. The obtained data were fitted with a four-parameter logistic regression. The inflection points of the curves can be used to compare attainable sensitivity. A.U. = arbitrary units.

### Recombinant ascorbate peroxidase fusion to anti-IgG nanobodies

Due to its stability and the breadth of its catalyzed colorimetric or chemiluminescent reactions that allow strong signal amplification, HRP is the most preferred enzyme for conjugation to secondary antibodies. However, it still has to be isolated from horseradish roots as a mixture of different isoforms, cannot be made in a practical scale and with a useful specific activity in *E. coli* (Krainer and Glieder, 2015), and it fails entirely as a genetic fusion to bacterially expressed nanobodies.

As an alternative, we tested the engineered APEX2 ascorbate peroxidase (Martell et al., 2012; Lam et al., 2015) as a fusion partner of the anti-mouse IgG1 Fc nanobody TP1107. The TP1107-APEX2 fusion was not only well-expressed and soluble in *E. coli* (Fig. S2 B), but also, it was active and efficiently catalyzed the oxidation of the initially colorless substrate Amplex Ultra Red to the highly fluorescent resorufin (Fig. 2 B). In line with previous reports (Lam et al., 2015), HRP seemed slightly more efficient than APEX2 in catalyzing this reaction. Nonetheless, low femtomole amounts of TP1107-APEX2 could be detected, suggesting its applicability e.g. in ELISA assays as well for immunohistochemistry and enzymatic antigen-localization in immunoelectron microscopy applications.

### Western Blotting with infrared fluorophore-linked anti-IgG nanobodies

A convenient alternative to peroxidase conjugation or fusion is the labeling of secondary antibodies with infrared fluorescent dyes. In fact, infrared fluorescent Western blotting has emerged as a superior alternative to classical ECL. It offers high signal-to-noise ratios, allows straightforward quantification due to signal linearity over many orders of magnitude and even enables the simultaneous dual color detection of multiple proteins. We thus labeled anti-IgG nanobodies site-specifically with the infrared fluorophore IRDye 800 at a C-terminal cysteine (Pleiner et al., 2015). The anti-rabbit IgG nanobody TP897 alone performed just as well as a commercial polyclonal anti-rabbit IgG secondary antibody, when it was used with rabbit polyclonal antibodies to detect various nucleoporins (Nups) in a *Xenopus* egg extract (Fig. 3 A). Similarly, the anti-mouse IgG1 Fc-specific nanobody TP1107 gave comparable or even higher signal intensities than a polyclonal anti-mouse IgG secondary antibody in Western Blotting on HeLa cell lysate (Fig. 3 B). Combinations of TP1107 with the compatible anti-mouse IgG1 Fab-specific nanobody TP886 or the anti-mouse kappa chain nanobody TP1170 provided a clearly better detection sensitivity than the polyclonal secondary antibody. TP1170 allows sensitive detection of IgG2a subclass mAbs, as shown here for the detection of the bacteriophage minor coat protein pIII (Fig. 3 C). We routinely found infrared fluorophore-labeled anti-IgG nanobodies to yield higher detection sensitivity than their HRP-conjugated counterparts. When combined with the compatible IRDye 680, dual color blots using e.g. mouse and rabbit primary antibodies are easily possible (not shown). In contrast to polyclonal secondary antibodies, IRDye-labeled anti-IgG nanobodies give also a clean and strong signal when pre-bound to primary antibodies before application. This makes a separate incubation with the secondary antibody dispensable and saves up to 2 h processing time per blot. We explored such a one-step staining strategy in more detail below for immunofluorescence.

**Figure 3.**
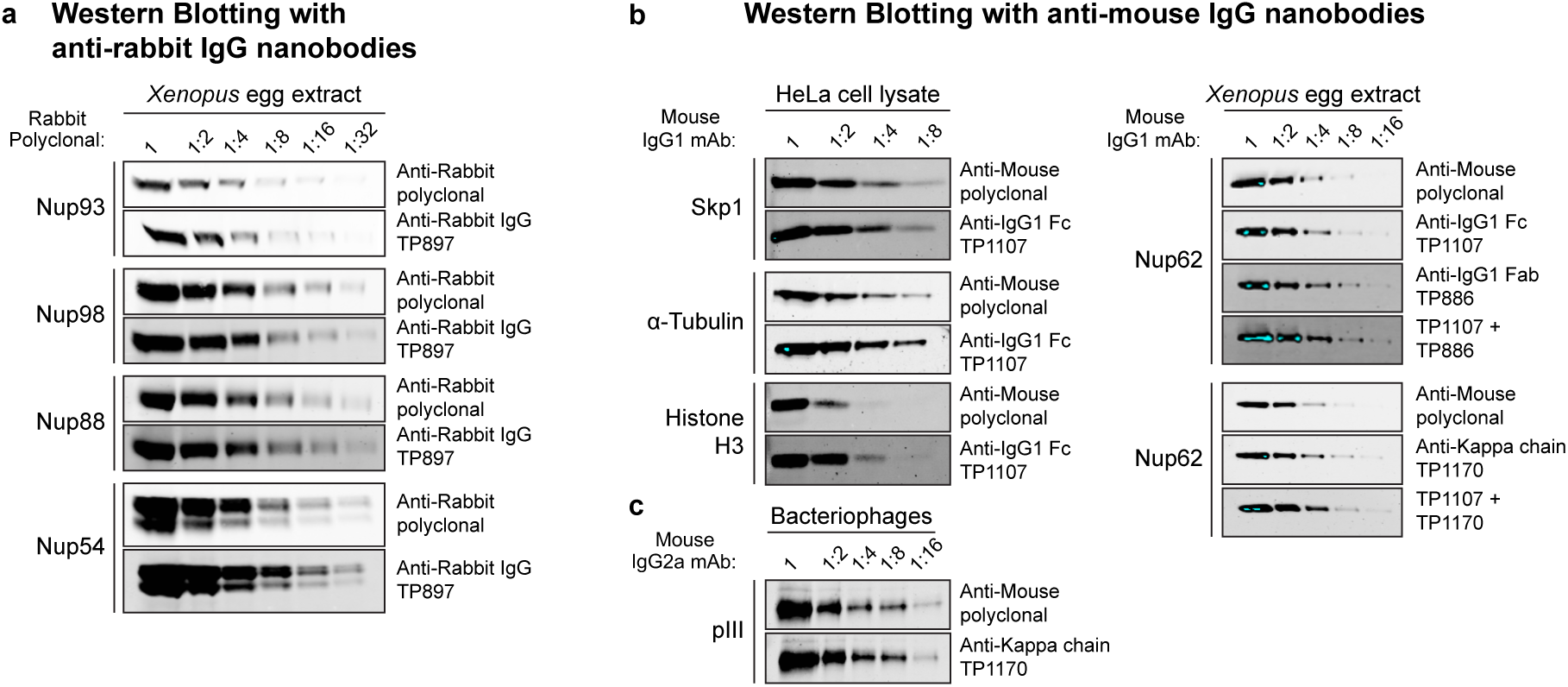
Western blotting with infrared dye labeled anti-IgG nanobodies. **(A)** A twofold dilution series of *Xenopus laevis* egg extract was analyzed by SDS-PAGE and Western Blotting. The indicated rabbit polyclonal antibodies were used to detect nucleoporins (Nups). These primary antibodies were then decorated either via IRDye 800-labeled goat anti-rabbit polyclonal IgG (1:5,000; LI-COR Biosciences, USA) or anti-rabbit IgG nanobody TP897 (10 nM). Blots were analyzed with an Odyssey Infrared Imaging System (LI-COR Biosciences, USA). **(B)** (Left panel) A twofold dilution series of HeLa cell lysate was analyzed by SDS-PAGE and Western Blotting. The indicated mouse IgG1 mAbs were decorated either via IRDye 800-labeled goat anti-mouse polyclonal IgG (1:1,340, 5 nM, LI-COR Biosciences, USA) or anti-mouse IgG1 Fc nanobody TP1107 (5 nM). (Right panel) A twofold dilution series of *Xenopus* egg extract was blotted and probed with anti-Nup62 mouse IgG1 mAb A225. It was then detected either via IRDye 800-labeled goat anti-mouse polyclonal IgG (5 nM), anti-mouse IgG1 Fc nanobody TP1107 (5 nM), anti-mouse IgG1 Fab nanobody TP886 (5 nM), anti-mouse kappa chain nanobody TP1170 (2.5 nM), a combination of TP1107 and TP886 or TP1107 and TP1170. Blue pixels indicate signal saturation. **(C)** A dilution series of filamentous bacteriophages was blotted and probed with an anti-minor coat protein pIII mouse IgG2a mAb. It was then decorated either via IRDye 800-labeled goat anti-mouse polyclonal IgG (2.5 nM) or anti-mouse kappa chain nanobody TP1170 (2.5 nM).

### Single and multi-color imaging with anti-IgG nanobodies

We next sought to assess the performance of the anti-IgG nanobodies as detection reagents in conventional indirect immunofluorescence. For this, cells are incubated sequentially with primary and secondary antibodies with intervening washing steps. Fluorophore-linked polyclonal secondary antibodies are routinely used for detection, since they can bind primary antibodies at multiple sites and thus deliver many fluorophores to enable large signal amplification. In contrast, individual anti-IgG nanobodies target only a single epitope per antibody (or two for symmetrical binding sites) and we therefore expected only modest signal amplification. Strikingly however, the anti-IgG1 nanobodies TP886 and TP1107, which specifically target IgG1 Fab and Fc fragment, respectively, not only performed well in Western Blotting, but also were well-behaved imaging reagents. For maximum brightness, we labeled these nanobodies with 2-3 fluorophores each at defined cysteines (Pleiner et al., 2015) and used them individually for the detection of mouse IgG1 mAbs in an indirect HeLa cell immunostaining (Fig. 4 A). Surprisingly, both were only slightly dimmer than the polyclonal mixture of anti-mouse secondary antibodies. We assume that the excellent nanobody signal is also due to less steric hindrance as compared to the much larger conventional secondary antibody. When both nanobodies were used in combination, we detected increased signal strengths that often were directly comparable to those obtained with the secondary antibody (e.g. for Vimentin or Ki-67) (see also Fig. S3 A). Importantly, despite a high labeling density with (the always somewhat sticky) fluorophores, we observed no detectable background staining with these anti-IgG nanobodies. This probably relates to the fact that the affinity of our nanobodies is very high, which allows their use at rather low nanomolar concentrations. The poor performance of the first anti-IgG nanobody generation indeed suggests that such excellent signal to noise ratio is not a trivial feature for a monovalent detection reagent.

**Figure 4.**
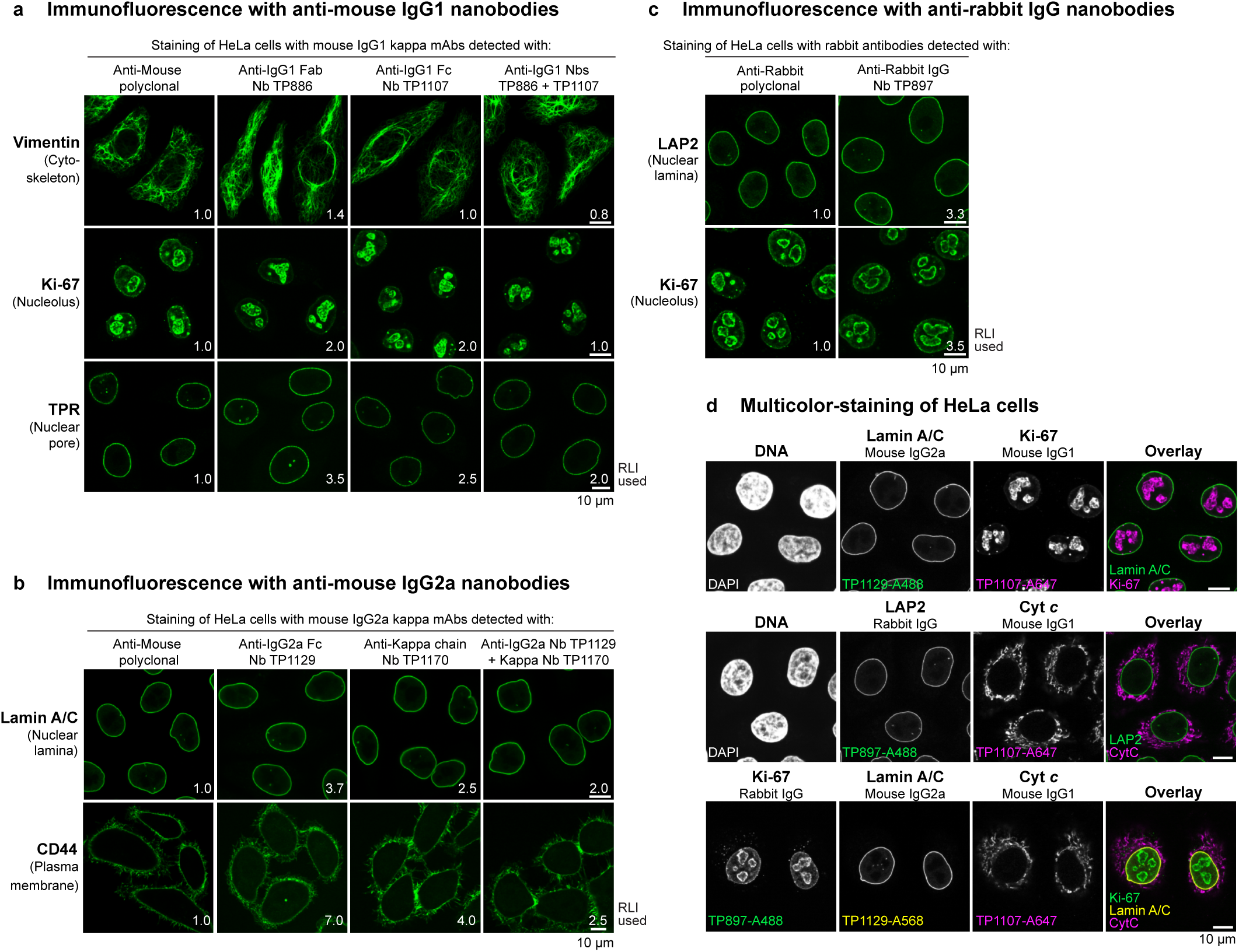
Imaging with anti-IgG nanobodies. **(A)** Immunofluorescence with anti-mouse IgG1 nanobodies. HeLa cells were stained with the indicated mouse IgG1 kappa mAbs. These primary antibodies were then detected with Alexa 488-labeled goat anti-mouse polyclonal antibody, anti-mouse IgG1 Fab nanobody TP886 or anti-mouse IgG1 Fc nanobody TP1107. A combination of TP886 and TP1107 yielded increased staining intensities. Laser intensities used to acquire the anti-IgG nanobody images were normalized to the intensity used to acquire the anti-mouse polyclonal antibody image (RLI = relative laser intensity is used here as a measure of fluorescence signal strength). **(B)** Immunofluorescence with anti-mouse IgG2a nanobodies. HeLa cells were stained with the indicated mouse IgG2a mAbs. These primary antibodies were then detected with Alexa 488-labeled goat anti-mouse polyclonal antibody, anti-mouse IgG2a Fc nanobody TP1129 or anti-kappa chain nanobody TP1170. A combination of TP1129 and TP1170 yielded increased staining intensities. **(C)** Immunofluorescence with anti-rabbit IgG nanobody TP897. HeLa cells were stained with the indicated rabbit antibodies. These primary antibodies were then detected with Alexa 488-labeled goat anti-rabbit polyclonal antibody or anti-rabbit IgG nanobody TP897. (**d**) Multicolor-staining of HeLa cells. HeLa cells were incubated with the indicated mouse IgG1, mouse IgG2a or rabbit IgG antibodies. These primary antibodies were detected via anti-mouse IgG1 Fc nanobody TP1107, anti-mouse IgG2a Fc nanobody TP1129 or anti-rabbit IgG nanobody TP897, respectively, labeled with the indicated Alexa dyes. The upper two panels show dual and the lower panel shows a triple co-localization.

For the detection of IgG2a subclass mAbs, we used a combination of two nanobodies, TP1129 and TP1170 (Fig. 4 B and Fig. S3 B). The IgG2a-specific nanobody TP1129 targets an epitope on the Fc-fragment and was obtained after affinity maturation of a lower affinity precursor (Fig. S3 C). Likewise, the kappa chain-specific nanobody TP1170 is an affinity-optimized variant, obtained after error-prone PCR, DNA shuffling and affinity selection (Fig. S3 D). TP1170 also proved effective in combination with the anti-IgG1 Fc nanobody TP1107 for the detection of IgG1 kappa mAbs (Figs. S3 E and F). The anti-rabbit IgG Fc nanobody TP897 can be used for the detection of polyclonal and monoclonal rabbit IgG (Fig. 4 C).

The presented nanobodies are specific for their respective IgG subclass, as shown in the specificity profiling dot blot assay (Fig. 1 B). We exploited this for multicolor imaging of HeLa cells with different IgG subclasses (Fig. 4 D). Mouse IgG1, mouse IgG2a and rabbit IgG-specific nanobodies did not show any crossreaction and consequently allowed for clean co-localization experiments. Even triple co-localizations were readily possible.

### Rapid one-step immunostaining and co-localization

The main reasons for separate incubation steps of primary and secondary IgGs in indirect immunofluorescence and Western blotting are the large size, as well as the bivalent and polyclonal nature of conventional secondary antibodies. If primary and secondary antibodies are pre-incubated, large oligomeric complexes form, which in immunofluorescence cannot easily penetrate into cells to reach their target and thus create background and poor signal (see Fig. 5 A). In contrast, anti-IgG nanobodies are monovalent and therefore do not crosslink primary antibodies. This allows streamlining the conventional immunostaining procedure to a single step. The primary antibodies are simply pre-incubated with fluorescently labeled anti-IgG nanobodies and then applied to cells together. After washing, the cells can be directly mounted for imaging. In such a workflow, anti-IgG nanobodies perform exceptionally well (Fig. 5 A). This time-saving protocol is also suitable for co-localization studies combining mouse and rabbit IgGs or combining mouse mAbs of different sub-classes. If the off-rate of the IgG pre-bound nanobodies were negligible over the staining period, then an exchange between the different pre-formed complexes would also be negligible. This would also make it unnecessary to use different IgG subclasses for multicolor imaging. We thus tested a multicolor staining workflow of HeLa cells relying solely on IgG1 subclass mAbs (Fig. 5 B). For this, we labeled anti-IgG1 Fc nanobody TP1107 with either Alexa 488, Alexa 568 or Alexa 647 maleimide and pre-incubated it with different IgG1 mAbs. The separately pre-incubated mixes were then combined and applied to HeLa cells for staining in one-step. Strikingly, we obtained clean dual and even triple co-localizations. In order to preclude an intermixing of colors, unlabeled TP1107 can be added in excess to the final mix and cells can be post-fixed after staining and washing.

**Figure 5.**
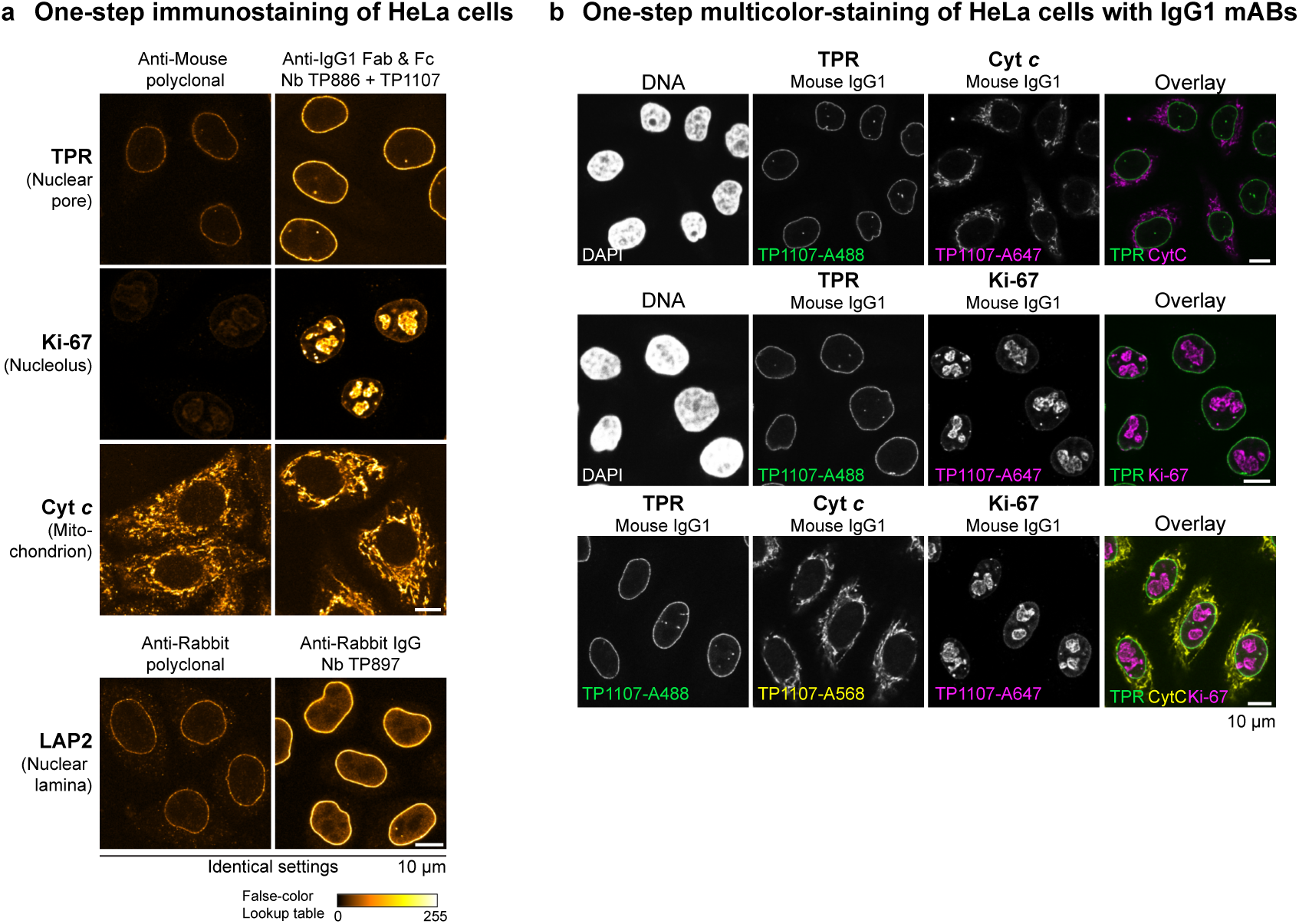
One-step immunostaining of HeLa cells with anti-IgG nanobodies. **(A)** The indicated mouse IgG1 mAbs were pre-incubated with an equal amount of Alexa 488-labeled goat anti-mouse secondary antibody or a combination of anti-mouse IgG1 Fab nanobody TP886 and anti-mouse IgG1 Fc nanobody TP1107. Likewise, the anti-LAP2 rabbit polyclonal antibody was pre-incubated either with Alexa 488-labeled goat anti-rabbit secondary antibody or anti-rabbit IgG nanobody TP897. The resulting mixes were then applied to fixed and blocked Hela cells. After washing, the cells were directly mounted for imaging. For every primary antibody, images were acquired under identical settings and pixel intensities are represented via a false-color lookup table. **(B)** Multicolor-staining of HeLa cells with mouse IgG1 subclass mAbs. The indicated mouse IgG1 mAbs were separately pre-incubated with Alexa 488, Alexa 568 or Alexa 647-coupled anti-mouse IgG1 Fc nanobody TP1107 and then mixed before staining HeLa cells in a single step. Washed cells were directly mounted for imaging.

### Super-resolution microscopy with anti-IgG nanobodies

Super resolution fluorescence imaging techniques offer the potential for observing sub-cellular structures at very small (e.g. nanometer) length scales (Huang et al., 2010; Sahl et al., 2017). However, these methods present new challenges for fluorescent labeling, because the spatial resolution of the images is comparable to the physical size of the probes. In the case of conventional antibodies, the antibody size is on the order of 10-15 nm, which may lead to a significant offset distance between the fluorophore and the epitope. At best, this offset may complicate the interpretation of super-resolution fluorescence image data, and in the worst case it makes it impossible to take full advantage of the increased resolution of the microscope.

Therefore, we reasoned that anti-Fab fragment or anti-kappa light chain nanobodies should be ideal imaging reagents for super-resolution microscopy, as they would enable small label displacement when used in conjunction with conventional primary antibodies. This would be essentially comparable to using directly labeled Fab fragments of primary antibodies, but does not require any extra work. In order to test this, we imaged microtubules of BS-C-1 cells by STORM with a resolution of approximately 20nm (Rust et al., 2006; Bates et al., 2007) (Fig. 6). Bound primary antibodies were detected either via Alexa 647-labeled polyclonal anti-mouse secondary antibodies or anti-mouse kappa chain nanobody TP1170. From the resulting images we selected straight regions of microtubule filaments and calculated the summed histogram of the localizations along the axis orthogonal to the filament axis. This yielded a measure of the filament width, which was obtained by fitting a Gaussian function to each histogram to determine the full width at half maximum (FWHM) of the cross-section. The distribution of filament widths measured for the two samples is shown in Fig. 6 B. In line with our initial expectations, we observed a striking difference in the microtubules’ apparent width for the two images. Microtubules stained via the polyclonal secondary antibody showed a median width of around 59.5 nm, which is in good agreement with electron microscopy studies of antibody coated microtubules (Weber et al., 1978) and previous STORM imaging (Bates et al., 2007). In contrast, staining with the anti-kappa chain nanobody yielded microtubules with a width of 37.5 nm, a remarkable ~22 nm reduction as a result of much lower label displacement as compared to the polyclonal secondary antibody. This result demonstrates, therefore, not only the significant offsets between epitope and fluorophore that may arise in conventional indirect immunostaining, but also the advantage of the smaller nanobody probe. Detection via the anti-kappa chain nanobody resulted in an image that more closely reflects the actual structure of the sample, suggesting its use as a superior secondary antibody for any super-resolution microscopy involving primary mouse antibodies.

**Figure 6.**
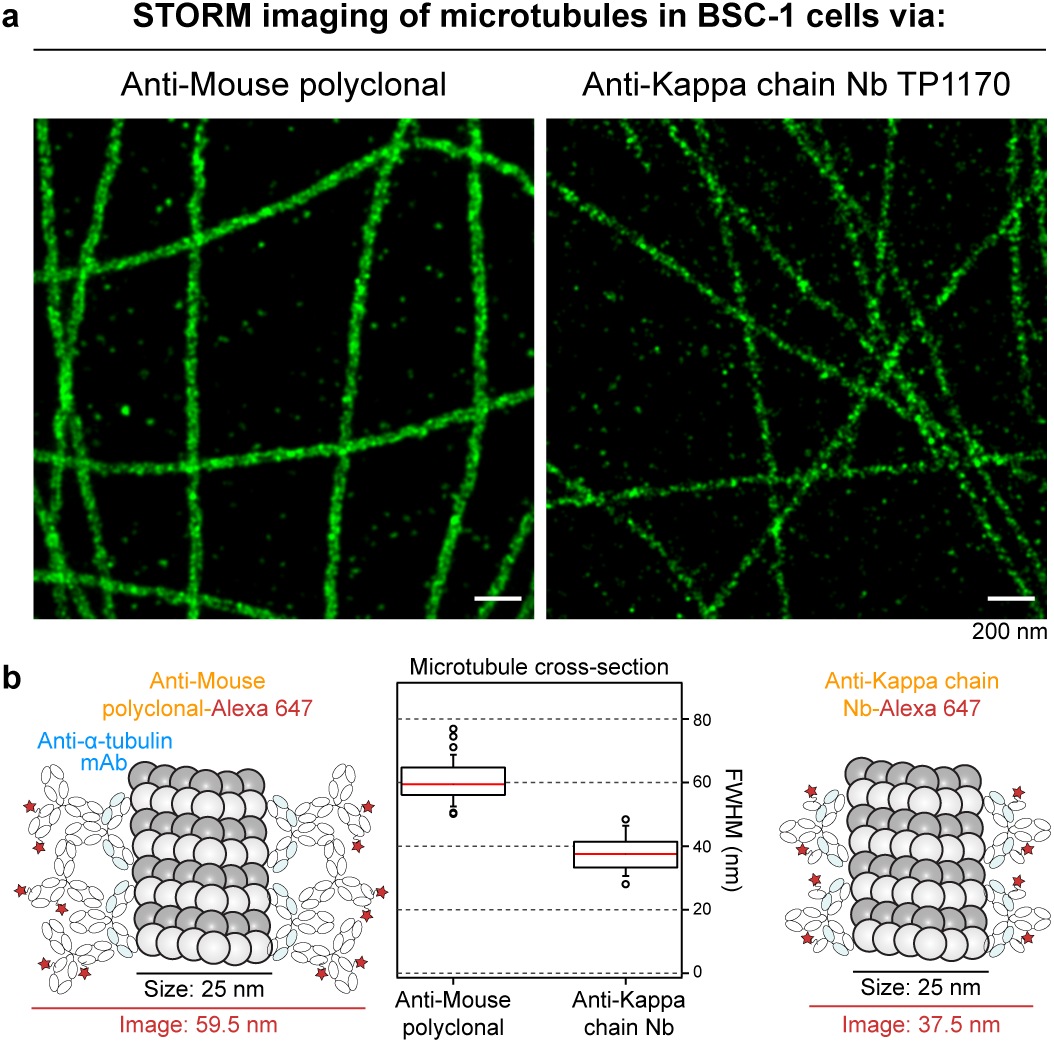
STORM imaging with anti-kappa chain nanobody TP1170. (**A**) BS-C-1 cells were stained with an anti-alpha tubulin monoclonal antibody (IgG1 kappa) and then detected with Alexa 647-labeled goat anti-mouse polyclonal antibody or Alexa 647-anti-mouse kappa chain nanobody TP1170. STORM images of the two samples show sub-diffraction limit organization of the tubulin filaments. (**B**) In order to quantify the effect of the label size on the apparent width of the filaments in the STORM images, averaged cross-sectional profiles of straight segments of filaments from the two samples were measured. First, the two labeling approaches are illustrated on the left and right of the figure, showing the expected smaller width for the nanobody labeling case. In the middle, box plots illustrate the results of the width analysis. In these measurements, the median width of the tubulin filaments decreased by a significant amount (from 59.5 nm to 37.5 nm) when stained with the anti-mouse kappa chain nanobody TP1170.

## Discussion

Due to the absence of more sustainable alternatives in the past, the great usefulness of polyclonal secondary antibodies in basic research certainly justified their animal-based production. However, in order to guarantee their constant supply to an ever-growing market, the producing companies had to dramatically increase their livestock, aim for very high antibody titres using aggressive hyper-immunization strategies causing strong side effects and increase the frequency and volume of collected bleedings. It is therefore not surprising that the global industrial scale production of antibodies causes severe animal welfare and ethical problems. The magnitude of these problems recently surfaced in the Santa Cruz Biotechnology scandal (Shen, 2013; Reardon, 2016).

Ideally, one should replace all animal immunization by selecting binders from synthetic libraries (Gray et al., 2016; Moutel et al., 2016; McMahon et al., 2017; Zimmermann et al., 2017). Yet, with a purely synthetic approach it is still not straightforward to obtain high-affinity binders. Further, the synthetic strategy is typically also inferior in terms of binder-specificity, because it lacks the stringent selection against self-reactivity that happens in antigen-exposed animals. The requirement for specificity is particularly high for secondary antibodies. We therefore see the here applied approach of using an immune library for binder selection as the best possible compromise. Since it is generally sufficient to obtain a few good nanobodies out of a small blood sample containing ~100 million lymphocytes, and since we found ways of further improving the initially found ones *in vitro*, there was no need for any hyper-immunization aiming at high titers. Importantly, once ideal nanobodies are identified, they are defined by their sequence and they can be renewably produced in *E. coli* at constant quality and without any further animal involvement. Since polyclonal secondary antibody production accounts for the largest share of immunized animals in the world, the anti-IgG nanobodies described in this study have the potential to make a great step forward towards reducing animal use and further contribute to a future of standardized recombinant antibodies (Marx, 2013; Bradbury and Plückthun, 2015a; Bradbury and Plückthun, 2015b).

We expect that our anti-IgG nanobodies will replace polyclonal secondary antibodies in many of their applications, e.g. in Western blotting and immunofluorescence. For both applications, their site-specific and quantitative modification with fluorophores via maleimide chemistry creates superior reagents with predictable label density and position. Furthermore, the precise targeting of primary mouse antibodies at the kappa chain with a specific nanobody can substantially reduce the label displacement in super-resolution microscopy. In the future, we will also explore the direct coupling of anti-IgG nanobodies with engineered cysteines onto colloidal gold particles for electron microscopy, which also suffers from the large linkage error introduced by bulky secondary antibodies.

Due to their monovalent and monoclonal nature, anti-IgG nanobodies do not crosslink primary antibodies and we exploited this for a one-step immunostaining workflow that saves valuable hands-on time and can also be extended to Western blotting. We envision that for routine stainings, preformed complexes of primary antibodies and labeled nanobodies can be prepared as stock solutions or simply bought from commercial suppliers. Due to the high affinity of the described nanobodies, the same strategy also enables multicolor immunostainings based on a single IgG subclass, which could also be relevant for flow cytometry sorting of specific cell types. This would be a cheaper and more flexible alternative to differentially labeled primary antibodies, it does not pose the risk of inactivating an antigen-binding site and it can easily be done if only small amounts of primary antibody are available.

Further, since the DNA sequences of these anti-IgG nanobodies are essentially synthetic building blocks, they can be genetically appended to the multitude of available tags, fluorescent proteins or enzymes to generate fusion proteins with novel functions for tailored applications in basic research and medical diagnostics, and also become valuable tools for immunology to study Fc or B cell receptors and downstream signaling cascades. Furthermore, anti-IgG nanobodies equipped with protease-cleavable affinity tags (Pleiner et al., 2015) will allow the native isolation of any antibody•target complex e.g. for structural studies by cryo-EM or functional assays.

Even though the here presented anti-IgG nanobody toolbox is already highly optimized, we will continue to extend it by identifying new nanobodies that decorate complementary binding sites and thus allow a further signal enhancement, and combine them with additional functional elements. In any case, it will be an open resource for all interested labs.

## Materials and methods

### Alpaca immunization

Two female alpacas, held at the Max Planck Institute for Biophysical Chemistry, were immunized 4 times with 1.0 mg polyclonal mouse or rabbit IgG at 3 week intervals. The anti IgG project turned out to be the so far most challenging nanobody project in the lab, because we aimed at an extremely low off-rate for imaging and blotting applications. We therefore resumed immunizations after a 12 months (rabbit IgG) or an 8 months break (mouse IgG). Nanobodies obtained after these late immunizations still showed very clear phage enrichment (> 1000-fold) even with femtomolar concentrations of the IgG baits. We therefore assume that they have very high affinity.

### Selection of anti-IgG nanobodies

The generation of nanobody immune libraries and the selection of antigen-specific nanobodies by phage display from these libraries were performed as previously described (Pleiner et al., 2015). IgG was biotinylated at accessible lysines by addition of a 4x molar excess of NHS-PEG_12_-biotin (Iris Biotech GmbH, Germany) for 2 h at room temperature in 1x PBS. Then the reaction was quenched and the excess of unreacted biotin separated from biotinylated IgG via buffer exchange into 50 mM Tris/HCl pH 7.5, 300 mM NaCl using PD-10 Desalting columns (GE Healthcare, USA).

### Expression and purification of untagged nanobodies

Bacterial expression plasmids for anti-IgG nanobodies will be distributed via Addgene. All nanobodies carry engineered cysteines and were expressed in the cytoplasm of *E. coli* NEB express F' (New England Biolabs, USA). A 50 ml preculture (2YT medium containing 50 µg/ml Kanamycin) was grown overnight at 28°C. The culture was then diluted with fresh medium to 250 ml. After 1 h of growth at 25°C, protein expression was induced for 3-5 h by adding 0.2 mM IPTG. After addition of 1 mM PMSF and 10 mM EDTA to the culture, bacteria were harvested by centrifugation, resuspended in lysis buffer (50 mM Tris/HCl pH 7.5, 300 mM NaCl, 10 mM imidazole, 5 mM DTT) and then lysed by sonication. The lysate was cleared by ultracentrifugation for 1.5 h (T647.5 rotor, Sorvall, 38,000 rpm) at 4°C. Nanobodies with engineered cysteines carried an N-terminal His_14_-*bd*NEDD8-tag and were affinity purified via Ni^2+^ chelate affinity chromatography. After washing with two column volumes (CV) of lysis buffer and one CV of maleimide-labeling buffer (100 mM potassium phosphate pH 7.5, 150 mM NaCl, 1 mM EDTA, 250 mM Sucrose), untagged nanobodies were eluted by on-column cleavage with 500 nM *bd*NEDP1 protease (Frey and Görlich, 2014) in maleimide-labeling buffer for 45 min at 4°C and labeled immediately with fluorophores. For longer storage, 10 mM DTT or TCEP were included in the maleimide-labeling buffer to keep cysteines reduced. Purified nanobodies were aliquoted and frozen in liquid nitrogen

### Site-specific fluorescent labeling of nanobodies with engineered cysteines

The fluorescent labeling of nanobodies with maleimide dyes was described in detail before (Pleiner et al., 2015). Briefly, stored nanobodies were thawed and the buffer was exchanged again to Maleimide-labeling buffer to remove the reducing agent, using either illustra NAP-5 or PD-10 desalting columns (GE Healthcare). For a standard labeling reaction, 5-10 nmoles of nanobody were rapidly mixed with 1.2x molar excess of fluorescent dye per cysteine on the nanobody and incubated for 1.5 h on ice. Free dye was separated from labeled nanobody by buffer exchange to Maleimide labeling buffer on illustra NAP-5 or PD-10 desalting columns. Quantitative labeling was quality controlled by calculating the degree of labeling (DOL). Fluorescently labeled nanobodies were always aliquoted, snap-frozen in liquid nitrogen and stored at −80°C until further use.

### Dot blot assay for anti-IgG nanobody specificity profiling

For profiling the binding of anti-IgG nanobodies to different IgG subclasses and to analyze their crossreaction to IgG from other species, a dot blot assay was performed. Nitrocellulose membrane was cut in strips and different IgGs (500 ng for polyclonal total IgG, Fab and Fc fragments; ~250 ng for monoclonal IgG in 1 µl) were spotted. Strips were blocked with 4 % milk (w/v) in 1xPBS for 30 min at room temperature. Then, nanobodies were added at ~300 nM in 1 ml milk for 30 min. After washing two times with 1x PBS for 10 min each, bound nanobodies were detected at 488 nm in a fluorescence scanner (Starion FLA-9000, Fujifilm, Japan). The following IgGs were used: IgG1 kappa mAb A225 (Cordes et al., 1995); IgG1 lambda (#010-001-331, Rockland, USA); IgG2a kappa (#02-6200, Thermo Fisher Scientific, USA); IgG2b kappa (#02-6300, Thermo Fisher Scientific, USA); IgG3 kappa (#401302, BioLegend, USA); polyclonal IgG Fab fragments (#010-0105, Rockland, USA); polyclonal IgG Fc fragments (#31205, Thermo Fisher Scientific, USA). Polyclonal IgG of the following species were used: rabbit (self-made, affinity-purified from serum); mouse (#I8765); rat (#I4131); goat (#I5256); sheep (#I5131); human (#I4506, all Sigma-Aldrich, USA) and guinea-pig (#CR4-10, Sino Biological, China).

### Native isolation of IgG with anti-IgG nanobodies

Polyclonal rabbit IgG from serum or mouse mAbs from hybridoma cell culture supernatant were isolated natively with anti-IgG nanobodies. For this, 0.3 nmoles of biotinylated nanobodies carrying a N-terminal His_14_-Biotin acceptor peptide-(GlySer)_9_-SUMOStar-(GlySer)_9_-tag were immobilized on 1 mg magnetic Dynabeads MyOne Streptavidin T1 (Thermo Fisher Scientific, USA). Excess biotin binding sites were quenched with biotin-PEG-COOH (#PEG1053, Iris Biotech, Germany). The beads were then incubated with 1 ml pre-cleared (10 min, 16,000g at 4°C) serum or hybridoma supernatant for 30 min at 4°C. After washing two times with wash buffer (50 mM Tris/HCl, 300 mM NaCl), nanobody-bound IgG was eluted by addition of 50 µl 0.5 µM SUMOStar protease (Liu et al., 2008) in wash buffer for 20 min on ice. An aliquot of the eluate was then analyzed by SDS-PAGE and Coomassie staining.

### Western Blotting

Bacteriophage protein III was detected with a mouse anti-pIII IgG2a mAb (#E8033S, New England Biolabs, USA). Mouse mAbs used for detection of human proteins in HeLa cell lysate were the following products: anti-Skp1 (clone H-6, #sc-5281, Santa Cruz Biotechnology, USA), anti-α-tubulin (clone DM1A, #T6199, Sigma-Aldrich, USA) and anti-Histone H3 (clone 96C10, #3638, Cell Signaling Technologies, USA). Polyclonal goat anti-mouse IgG coupled to IRDye 800CW (#925-32210; LI-COR Biosciences, USA) was used to detect primary mouse antibodies at a dilution of 1:1340 (5 nM). Polyclonal rabbit antibodies against *Xenopus laevis* nucleoporins Nup98, Nup93, Nup54 and Nup88 were prepared in the lab (Hülsmann et al., 2012). Polyclonal goat anti-rabbit IgG coupled to IRDye 800CW (#925-32211; LI-COR Biosciences, USA) was used to detect primary rabbit antibodies at the lowest suggested dilution of 1:5,000. Anti-mouse IgG1 Fab nanobody TP886 (5 nM), anti-mouse IgG1 Fc nanobody TP1107 (5 nM) and anti-rabbit IgG nanobody TP897 (10 nM) were labeled with a single IRDye 800CW maleimide (#929-80020, LI-COR Biosciences, USA) via a C-terminal cysteine and used at the indicated concentrations in 4 % (w/v) milk in 1x PBS. Polyclonal goat anti-mouse-HRP conjugate was from DakoCytomation (Denmark) and used at 1:1,000 dilution (5 nM). Anti-mouse IgG1 Fc nanobody TP1107 was conjugated to maleimide-activated HRP (#31485, Thermo Fisher Scientific, USA) via a C-terminal cysteine by mixing both in equimolar amounts and incubation for 1h at room temperature. The conjugate was used at 5 nM in 4 % (w/v) milk in 1x PBS. The ECL solution was self-made and contained 5 mM Luminol (#A4685, Sigma-Aldrich, USA), 0.81 mM 4-Iodophenylboronic acid (#471933, Sigma-Aldrich, USA) and 5 mM freshly added H_2_O_2_ in 0.1 M Tris/HCl pH 8.8.

### Amplex Ultra Red assay

APEX2 was derived from pTRC-APEX2 (Addgene plasmid #72558), which was a gift from Alice Y. Ting (Lam et al., 2015). The anti-mouse IgG1 Fc nanobody TP1107-APEX2 fusion was expressed from pTP1135 with an N-terminal His_14_-*bd*NEDD8-tag in *E. coli* NEB express F’ (New England Biolabs, USA) for 6 h at 25°C in the presence of 1 mM of the heme precursor 5-aminolevulinic acid (#A3785, Sigma-Aldrich, USA). Following lysis, the protein was purified by nickel chelate affinity chromatography and eluted by cleavage with 500 nM *bd*NEDP1 protease (Frey and Görlich, 2014) in 100 mM potassium phosphate pH 7.5, 150 mM NaCl, 250 mM sucrose. The final assay mix contained 160 µM Amplex Ultra Red, 160 µM H_2_O_2_ in either 100 mM Citrate pH 6.6, 150 mM NaCl (optimal pH for APEX2) or 100 mM potassium phosphate pH 6.0, 150 mM NaCl (optimal pH for HRP). 50 µl of this mix was used per reaction. Anti-mouse IgG1 Fc nanobody TP1107-APEX2 was titrated from 167 nM to 470 fM in a 1.8-fold dilution series and 2 µl of each dilution added to 50 µl reaction mix in triplicates. HRP (#31490, Thermo Scientific, USA) was titrated from 31 nM to 5 fM in a 2.4-fold dilution series and 2 µl per dilution added to 50 µl reaction mix in triplicates. The 96-well plate containing these reactions was incubated at room temperature for 30 min and then resorufin fluorescence was measured at 590 nm (530 nm excitation) in a Bio-Tek Synergy HT Multi-Detection Microplate Reader (BioTek Instruments Inc., USA).

### Immunofluorescence

HeLa cells grown on glass coverslips were fixed for 10 min at room temperature with 3 % (w/v) paraformaldehyde (PFA) and then washed two times with 1x PBS for 5 min each. Residual PFA was quenched by incubation with 50 mM NH_4_Cl in 1x PBS for 5 min. After two washes with 1x PBS for 5 min each, the cells were permeabilized with 0.3 % (v/v) Triton-X-100 for 3 min. Then the cells were washed three times quickly with 1x PBS and blocked for 30 min with 1 % (w/v) BSA in 1x PBS (blocking buffer). Following blocking, the coverslips were stained with primary antibody, which was diluted in blocking buffer, in a humid chamber for 1 h at room temperature. The coverslips were then washed two times in 1x PBS for 15 min each and added again to a humid chamber for incubation with secondary antibody or anti-IgG nanobody diluted in blocking buffer. Afterwards, the cells were washed two times in 1x PBS for 15 min each and the coverslips mounted with Slow Fade Gold (Thermo Fisher Scientific, USA) for imaging on a Leica TCS SP5 confocal microscope equipped with hybrid detectors (Leica, Germany).

For methanol fixation, the cells were incubated with −20°C-cooled methanol for 6 min at room temperature, washed two times in 1x PBS for 5 min each and then blocked in blocking buffer. The staining was performed as described above.

### Antibodies for immunofluorescence

The following rabbit antibodies were used for immunofluorescence on HeLa cells: anti-Lap2 polyclonal antibody (1:100 dilution, #14651-1-AP, Proteintech, UK); anti-Ki-67 mAb clone D3B5 (1:200 dilution, #9129, Cell Signaling Technologies, USA). The following mouse mAbs were used for immunofluorescence on HeLa cells: anti-Vimentin mAb clone V9 (1:10 dilution of Hybridoma supernatant, kind gift of Mary Osborn); anti-Ki-67 mAb clone B56 (1:50 dilution, #556003, BD Bioscience, USA); anti-TPR mAb 203-37 (1:500 dilution, Matritech Inc., USA; (Cordes et al., 1997)); anti-Cytochrome (Cyt) c mAb clone 6H2.B4 (1:50 dilution, #556432, BD Bioscience, USA); anti-Lamin A/C mAb clone 4C11 (1:50 dilution, #4777T, Cell Signaling Technologies, USA); anti-CD44 mAb clone 156-3C11 (1:200 dilution, #3570T, Cell Signaling Technologies, USA). Polyclonal goat anti-rabbit IgG (#111-545-003) and goat anti-mouse IgG (#115-545-003, Jackson ImmunoResearch, USA) coupled to Alexa Fluor 488 were used as secondary antibodies at 1:150 dilution (~33 nM). Anti-IgG nanobodies were labeled with maleimide Alexa Fluor dyes at engineered surface cysteines (Pleiner et al., 2015) and used at 20 nM. The used nanobodies had the following degree of labeling: TP886-Alexa 488 = 1.9, TP1107-Alexa 488 = 2.7, TP1107-Alexa 647 = 2.2, TP1129-Alexa 488 = 2.5, TP1129-Alexa 568 = 2.0, TP1079-Alexa 488 = 2.2, TP897-Alexa 488 = 2.2.

### STORM imaging of microtubules in BS-C-1 cells

BS-C-1 cells were stained with an anti-alpha tubulin monoclonal antibody (1:200 dilution, # T6074, Sigma Aldrich, USA) after PFA fixation as described above for HeLa cells. STORM imaging was carried out using a custom built microscope, similar to what has been described previously (Bates et al., 2012). Briefly, 642nm laser light was used to illuminate the sample, and fluorescence was detected with an EMCCD camera (Andor Ixon DU860), after filtering with a bandpass filter (ET700/75, Chroma Technologies). Raw STORM data was analyzed with custom written software, and STORM images of each sample were rendered using summed Gaussian functions. For calculation of the cross-section histograms, multiple straight segments of tubulin filaments were selected from the STORM images. For each straight filament segment, a line was overlaid the segment in order to define the filament axis. Next, a set of rectangular regions of interest (ROIs) was created, aligned with the segment, spanning the cross-section of the filament. The ROI length was set equal to the segment length, and a user-selectable ROI width, which was chosen to be 5 nm for this analysis (the bin width). By counting the number of localizations falling within each ROI, a histogram corresponding to the cross-sectional profile of the STORM image of a filament, averaged along the segment length, was generated. To measure the width of the cross-section, a Gaussian function was fit to the histogram, and the full width at half maximum was calculated. The distribution of measured filament widths is shown in Fig. 6 B.

## Acknowledgements

We would like to thank Ulrike Teichmann and Rolf Rümenapf for alpaca care and immunization; Jens Krull, Heinz-Jürgen Dehne, Gabriele Hawlitscheck and Tanja Gilat for excellent technical assistance; Philip Gunkel, Volker Cordes, and Mary Osborn for advice and antibodies; Trevor Huyton for critical reading of the manuscript and the Max-Planck-Gesellschaft as well as the Deutsche Forschungsgemeinschaft (DFG) (SFB1190) for funding this work.

## Author contributions

TP: Conception and design, Acquisition of data, Analysis and interpretation of data, Drafting and revising the article; MB: Acquisition of data, Analysis and interpretation of data, Drafting and revising the article; DG: Conception and design, Analysis and interpretation of data, Drafting and revising the article.

---

**Figure S1.**
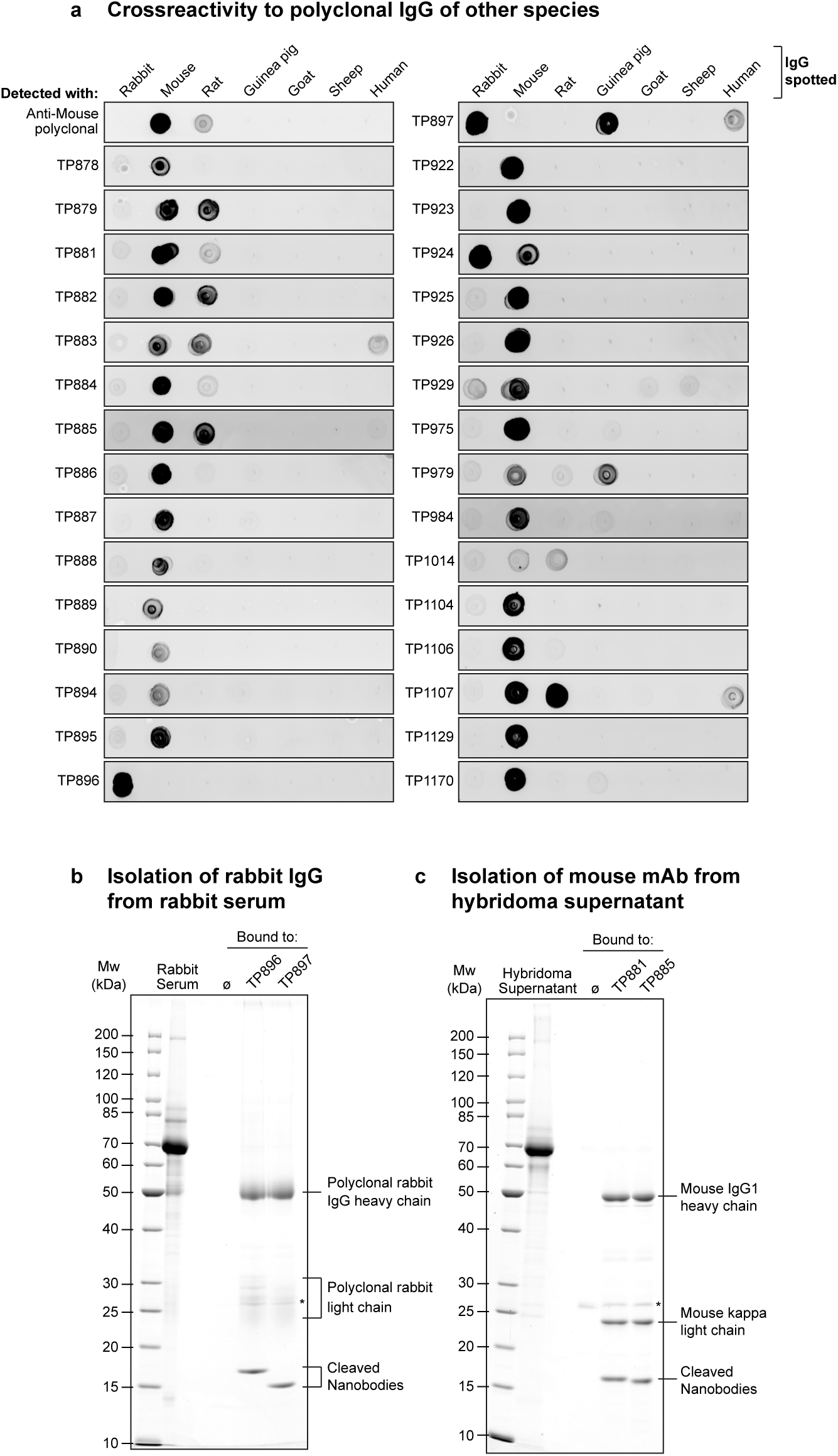
Species crossreactivity profiling and native target IgG isolation. **(A)** Crossreactivity profiling of all anti-IgG nanobodies. Using the same Dot blot assay as described in Fig. 1 B, the crossreactivity of anti-IgG nanobodies to polyclonal IgG from the indicated species was determined. **(B)** Isolation of polyclonal rabbit IgG from rabbit serum. Anti-rabbit IgG nanobodies TP896 and TP897 carrying an N-terminal Avi-SUMOStar tag were biotinylated and immobilized on magnetic Streptavidin beads. After incubation with crude rabbit serum and washing, nanobody-bound polyclonal rabbit IgG was specifically eluted via SUMOStar protease cleavage in physiological buffer. Empty beads served as negative control. **(C)** Isolation of anti-Nup62 mouse IgG1 kappa mAb A225 from hybridoma supernatant with anti-mouse IgG1 nanobodies TP881 and TP885 as described in **B**. The asterisk indicates the SUMOStar protease used for elution.

**Figure S2.**
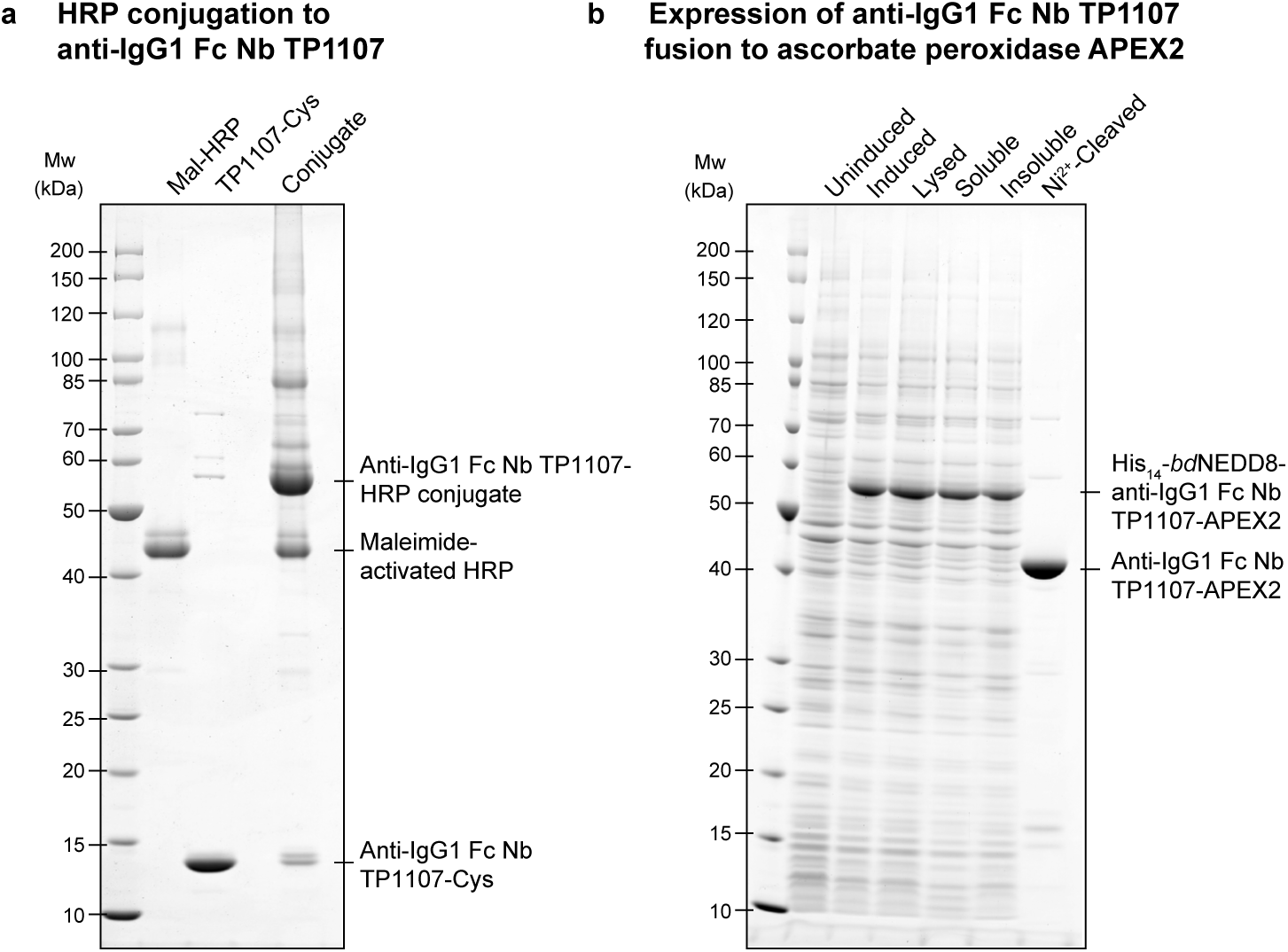
Anti-IgG nanobody conjugation to HRP and fusion to APEX2. **(A)** Anti-mouse IgG1 Fc nanobody TP1107 with a C-terminal cysteine was conjugated to maleimide-activated horseradish peroxidase (HRP) by incubation of equimolar amounts for 1 h at room temperature. **(B)** Expression of anti-mouse IgG1 Fc nanobody TP1107-ascorbate peroxidase (APEX2) fusion in *E. coli*. After binding to nickel beads via the N-terminal His_14_-*bd*NEDD8-tag, untagged fusion protein was eluted by on-column *bd*NEDP1 cleavage (Frey and Görlich, 2014).

**Figure S3.**
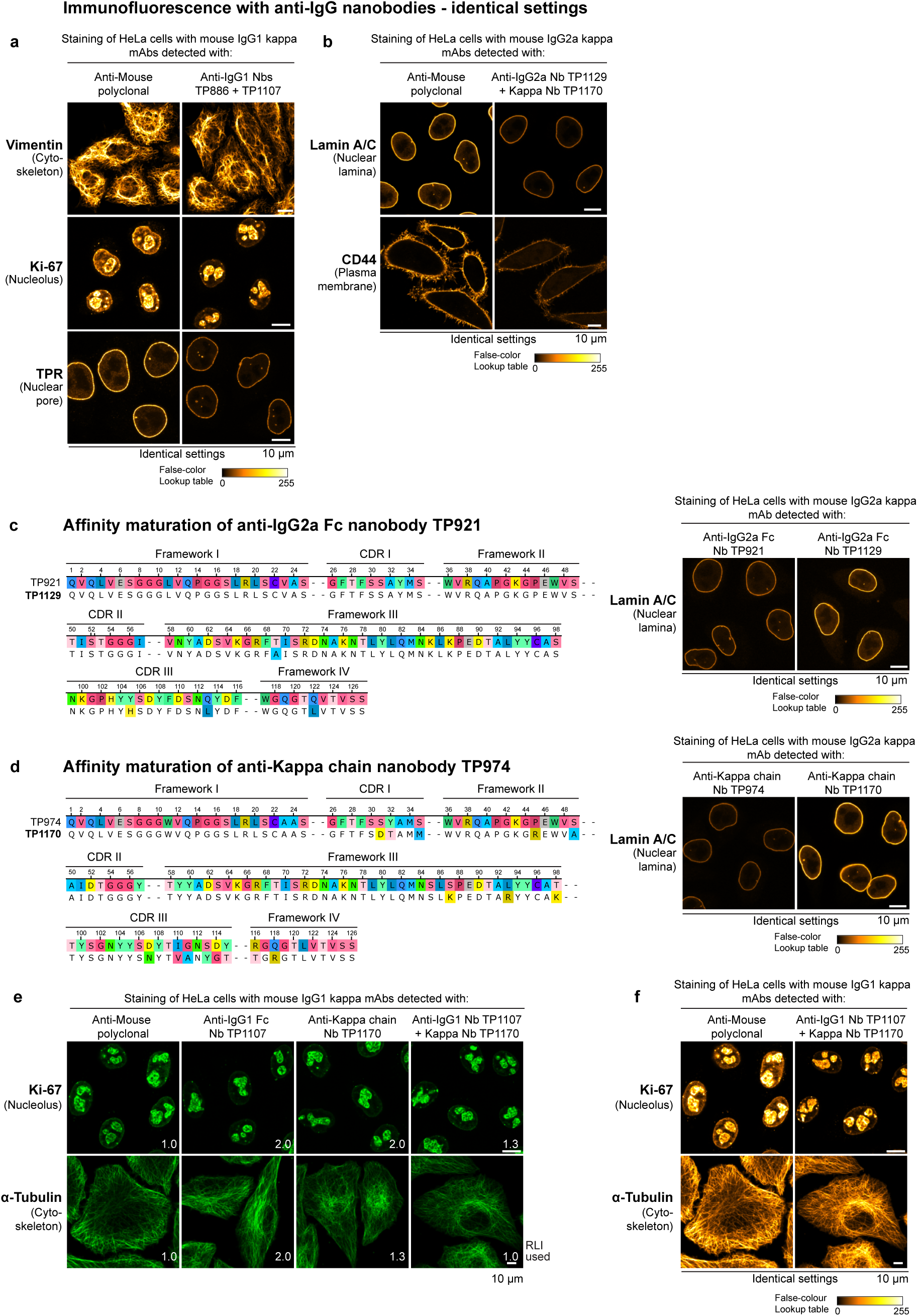
Immunofluorescence with anti-mouse IgG nanobodies. **(A-B)** Images for a given mAb or polyclonal antibody were acquired under identical settings and pixel intensities are represented via a false-color lookup table. **(A)** HeLa cells were stained with the indicated mouse IgG1 mAbs. These primary antibodies were then detected with Alexa 488-labeled goat anti-mouse polyclonal antibody or a combination of anti-mouse IgG1 Fab nanobody TP886 and anti-mouse IgG1 Fc nanobody TP1107. **(B)** HeLa cells were stained with the indicated mouse IgG2a mAbs. These primary antibodies were then detected with Alexa 488-labeled goat anti-mouse polyclonal antibody or a combination of anti-mouse IgG2a Fc nanobody TP1129 and anti-kappa chain nanobody TP1170. **(C)** Protein sequence alignment of anti-mouse IgG2a nanobody TP921 and the variant TP1129 obtained after affinity maturation. HeLa cells were stained with a mouse IgG2a mAb targeting Lamin A/C. The mAb was detected via TP921 or TP1129 labeled with a single Alexa 488 dye and the images acquired under identical settings. **(D)** Protein sequence alignment of anti-mouse kappa chain nanobody TP974 and the variant TP1170 obtained after DNA shuffling and affinity maturation. HeLa cells were stained with a mouse IgG2a mAb targeting Lamin A/C. The mAb was detected via TP974 or TP1170, both labeled with two Alexa 488 dyes. **(E)** HeLa cells were stained with the indicated mouse IgG1 kappa mAbs. These primary antibodies were then detected with Alexa 647-labeled goat anti-mouse polyclonal antibody, anti-mouse IgG1 Fc nanobody TP1107 or anti-mouse kappa chain nanobody TP1170. A combination of TP1107 and TP1170 yielded increased staining intensities, see **(F)** for identical settings scan. RLI = relative laser intensity (as defined in Fig. 4 A).

**Supplementary Table 1.**
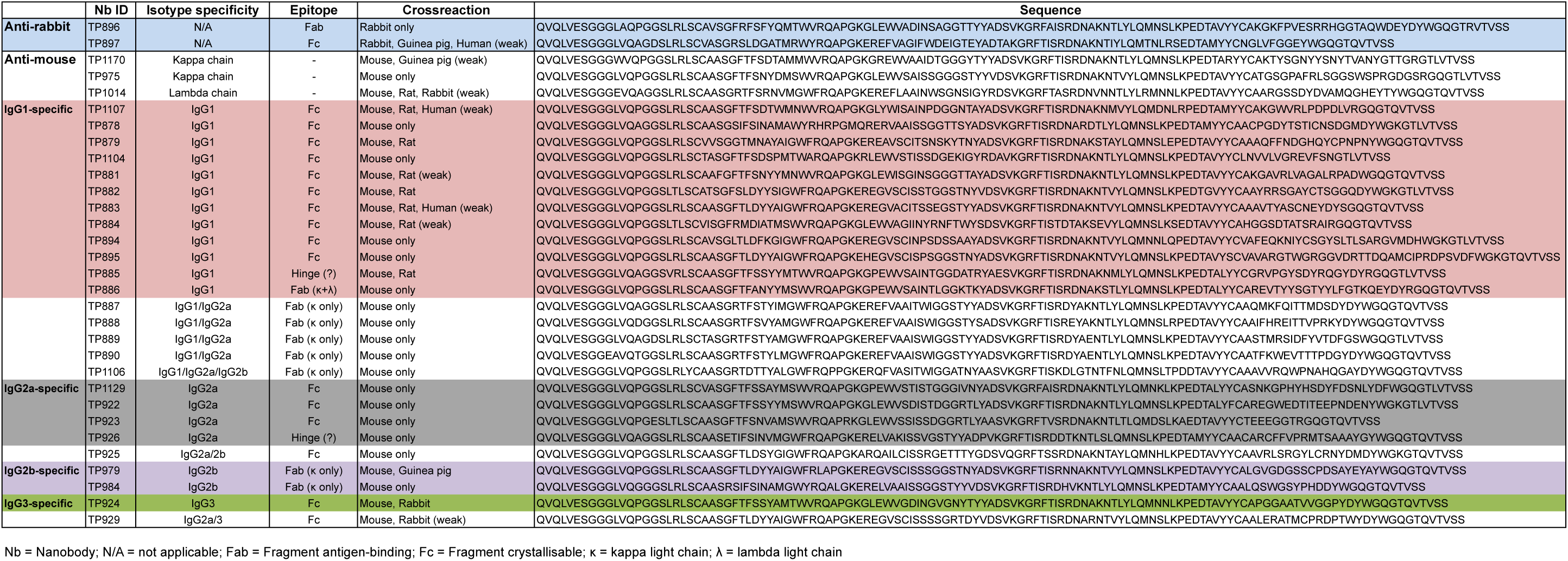
Anti-IgG nanobody protein sequences

